# Geometric Evaluation and Optimization of Maize Leaf Morphology and Canopy Architecture

**DOI:** 10.1101/2025.08.01.668074

**Authors:** Zhaocheng Xiang, Yufeng Ge

## Abstract

Maize leaf and canopy are the primary media to interact with environmental factors, including light interception, water evaporation, wind fluctuation, etc. Leaf morphology and canopy architecture affect the interaction process and activity significantly. Maize canopy architecture, a critical determinant of light capture and photosynthetic efficiency, is influenced by complex interactions between leaf morphology and arrangement. This study aimed to explore the relationships between key leaf morphological traits (midrib shape, blade contour, undulations) and canopy architecture in maize, and to identify optimal leaf morphologies for light interception. We employed correlation analysis and stepwise Bayesian optimization to quantify the relationships between leaf traits and to identify optimal leaf morphologies for maximizing leaf area and minimizing self-shading. Our results revealed that midrib shape and leaf width distribution positively influence leaf area, while undulation has a bidirectional effect depending on its orientation. Optimization results showed that optimal canopies feature upright top leaves, intermediately erect middle leaves, and relatively flat bottom leaves. This study provides a quantitative framework for understanding and predicting optimal canopy structure in maize, which can inform breeding strategies aimed at improving light utilization and yield potential.

## 1. Introduction

Maize (*Zea mays*), one of the major field crops, plays an important role in human society. It is widely planted all over the world across various climate regions. The key maize indicators, including maize planting area, yield and production, have been increased significantly after the Green Revolution in 1960s (FAO, 2024). The planting area increased two times, and the average yield increased three times. Consequently, global maize production increased about six times. There are several reasons that drove the increase in global maize production. The first and the most important factor is the increase in population which increased the demand for maize as food and energy source. Maize is the primary direct food source in many Latin American, Sub-Sahara African and Asian countries (Erenstein et al., 2022). Maize also contributes to human food and nutrition indirectly serving as animal feeds, sugar and starchy feedstock. Furthermore, maize is utilized as biofuel by fermentation processes which has been massively commercialized in United States, Brazil and Europe (Mussatto et al., 2010). The second factor is the development of agricultural and biological technologies. The improvement in management practices, such as irrigation, fertilization and pest management, has increased resource use efficiency in agricultural production and improved its quantity and quality (Rudel et al., 2009; Tilman et al., 2002). The essence of biological improvement is the breeding technique which developed the modern varieties of many crops, including maize (Duvick, 2005; Evenson & Gollin, 2003). Genetic engineering has boosted modern maize breeding because of its efficiency compared to traditional breeding (Andorf et al., 2019). Identification of the traits and loci is needed to edit the target gene regions and acquire the interested traits. Plant phenotyping and quantitative genetics are the main tools to build the linkage between interested traits and target genes. Maize leaf is the primary organ for light interception, carbon assimilation, photosynthesis and respiration. The morphological traits on maize leaf can indicate plant health and performance comprehensively. However, the quantitative description of maize leaf morphology is still an open question. This problem hinders the implementation of high throughput phenotyping at the leaf level and the genetic investigation about leaf traits. Furthermore, the mathematical and geometric relations between morphological traits on leaf are poorly studied which provide inadequate guidance for ideotype design.

Currently, plant phenotyping has been implemented on simple maize leaf morphological traits, including leaf length (Daviet et al., 2022; Zhang et al., 2017), width (Y. Wang et al., 2019), area (Thapa et al., 2018), angle (Bao et al., 2019; He et al., 2024), curvature (Liu et al., 2024; Y. Wang et al., 2019), rolling (Baret et al., 2018), twisting (Wu et al., 2022), etc. However, those simple leaf traits provide insufficient interpretation on maize growth status and physiological performance. Every individual maize is a complex biological system. The dim light shed on those simple traits could not be evident enough to build the network of interactions between genotype, environment and management practices. Systematic modeling of maize leaf geometry is a key step for quantification of the leaf morphology and phenotyping on the subtle leaf features. Several maize models have been proposed to describe the whole plant profile and used in simulation. ADEL-maize is the first L-system model for maize (Fournier & Andrieu, 1999). It used iterative phytomer to model the repeating elements in maize, which is composed of internode and leaf. GREENLAB implemented multi-fitting method to optimize maize growth parameters and visualized the outputs in 3D (Guo et al., 2006; X. Wang et al., 2024). GRAAL-CN integrated plant morphogenesis and carbon and nitrogen assimilation to model plant growth during the vegetative stages (Drouet & Pagès, 2007). However, the maize models address the importance and consistency of the overall maize plant instead of the local characteristics on the specific organs. The subtle features on maize leaf are investigated limitedly.

Compared to leaves from other plants, maize leaf is long and slim whose shape is like a dagger knife. The shape of maize leaf is simple and linear as its margin doesn’t have teeth or serration at the edge. The venation is also simple, which only has parallel veins. The entire leaf surface is also smooth without breaking points or lines. However, maize leaf is still geometrically complex in 3D space. The curves on maize leaf contribute to the geometric complexity overwhelmingly. There are three primary curves on maize leaf: midrib, cross section and blade contour. Midrib is like a conic curve due to gravity effects. The ascending part of midrib is strong which can support the whole leaf weight while the descending part is thin and dangling. Some studies utilized parabola to fit midrib shape (Ledent, 2006; Qian et al., 2023). However, maize midrib is not an exact parabola due to the different morphology of the ascending and descending parts (Ford et al., 2008). Prévot et al. (1991) facilitated two parabolas to fit midrib which are connected at the highest point. The method can fit midrib shape on a broad basis that is also implemented in ADEL-Maize model. But heavy computation is necessary to ensure the insertion point is geometrically smooth. Furthermore, midrib twisting also increases the geometric complexity since it transforms the midrib curve from 2D plane to 3D space. The cross section of maize leaf is like a gull wing. Traditionally, the traits related to cross section were simplified into leaf width. Wen et al. (2024) proposed blade included angle to represent the geometry of maize leaf midrib and laminas. The blade contour is the projection of the leaf. It is determined by leaf width distribution. Sanderson et al. (1981) proposed a polynomial trigonometric function to fit the width distribution. However, those descriptions of maize leaf elements are discrete which are insufficient to evaluate maize leaf morphology systematically.

Canopy architecture plays an important role in light interception. Canopy is composed of leaves that can be divided into three groups: top layer, middle layer and bottom layer. Therefore, the leaf shape at different layers affects the overall canopy architecture and light interception ability. Tian et al. (2024) proposed a smart canopy for maize which has upright top leaves, less erect middle leaves and relatively flat bottom leaves. Light within maize canopy can be more evenly distributed and improve radiation use efficiency. However, the description is empirical and qualitative. Ideotype design and modern breeding for maize require quantification of canopy architecture. However, it is still an open question due to the high geometrical and morphological complexity and diversity.

The main contribution of this paper is a quantitative evaluation on maize leaf morphology and optimization for leaf area and canopy architecture based on the model proposed by Xiang and Ge (2025). We used the parametric model to analyze the correlation between maize leaf morphological characteristics and leaf area indicators. We proposed stepwise Bayesian method to optimize leaf area under given configuration of fixed leaf length and width. We implemented multi-objective Bayesian optimization to study the geometric characteristics of leaves at the top, medium and bottom layers to describe the canopy architecture for light interception.

## 2. Materials and Methods

### 2.1 3D Leaf Generation and Morphological Characteristics

The 3D leaf generation was built upon the maize leaf modeling method proposed by Xiang and Ge (2025). The method decomposes a maize leaf into midrib, cross section and blade contour and then triangle mesh is created by translation, rotation and scaling of the three elements. We modified the original model to explore more complicated maize leaf morphological characteristics.

#### 2.1.1 Midrib Curvature and Aspect Ratio

The midrib curvature is represented by a hyperbolic spiral which has parametric equations for the Cartesian coordinate as Eq. 1. The intermediate parameter, *ϕ*, was configured by constant left boundary, 3.0, and variable right boundary between 3.6 and 5.5. If curvature has the unique extreme when *ϕ* is 4.49 where the curvature increases in the left and decreases in the right. The midrib curvature is controlled by three shape parameters, including M1, M2 and the right boundary. However, if the right boundary of *ϕ* and leaf length are fixed, two midrib curvatures have the same shape when the ratios between M1 and M2 are equal. Therefore, we simplified the midrib representation into the shape ratio and the right boundary. Consequently, M2 is always 1 and M1 is the shape ratio in Eq.1. The range of the shape ratio is between 0.2 and 5 since the M1 and M2 are within 1 and 5.

Midrib curvature was represented using a hyperbolic spiral, defined by the parametric equations in Eq. 1. The intermediate parameter, *ϕ*, was configured with a constant left boundary of 3.0 and a variable right boundary ranging from 3.6 to 5.5. A unique curvature extremum occurs at *ϕ* = 4.49, with curvature increasing to the left and decreasing to the right of this point. While midrib curvature is influenced by three shape parameters (M1, M2, and the right boundary of *ϕ*), it was observed that, for a fixed right boundary and leaf length, curvature shapes are identical when the ratio between M1 and M2 is constant. This allowed for a simplification of the curvature representation, using the shape ratio (*sr* = *M*1/*M*2) and the right boundary of *ϕ*. Therefore, M2 was fixed at 1, and M1 in Eq. 1 represents this shape ratio, ranging from 0.2 to 5, based on the initial constraints of M1 and M2 being within 1 and 5.

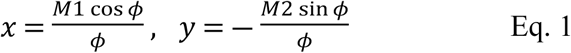

To characterize the overall midrib curvature, the aspect ratio was utilized as an alternative indicator for leaf elevation angle. This aspect ratio was calculated based on the elevation and elongation of the midrib. Elevation was defined as the vertical distance between the highest and lowest points of the midrib, while elongation represented the horizontal distance between its leftmost and rightmost points.

#### 2.1.2 Projection Area and Rugosity

The projection area was calculated by projecting the 3D mesh onto the XOY plane and calculating the area of the resulting polygon. Rugosity was introduced by Risk (1972) to evaluate the surface complexity of coral reef. Rugosity is defined as the ratio of a straight-line distance to the actual distance along a surface, as determined by a flexible chain. Leaf rugosity is the ratio of leaf area to projection area. Midrib curvature, self-twisting and undulation influence rugosity. Specifically, erect leaves, characterized by a smaller projection area, exhibit a proportionally higher rugosity. The rugosity metric can be used to evaluate leaf characteristics related to shading on lower leaves and the evenness of light distribution within canopy.

#### 2.1.3 Self-Twisting

To simulate the natural self-twisting observed in maize leaves, a twisting angle was implemented that varied linearly along the midrib from leaf base to tip. This was achieved by progressively rotating the midrib itself along its longitudinal axis. The degree of rotation at any point along the midrib was determined by a linear function, with the angle increasing proportionally to the distance from the base. This approach allowed for the recreation of a range of twisting phenotypes, from minimal twisting to pronounced helical configurations, capturing the natural variation observed in maize leaves. The twisting angle at the leaf tip was defined as a user-input parameter, enabling precise control over the degree of self-twisting in the model.

#### 2.1.4 Undulation

Undulation represents the wave-like patterns along the leaf edge and surface. The frequency and amplitude of undulation were varied to simulate different undulation phenotypes. Undulation was generated by the rotation of the cross-section frames around two directions during leaf generation. One direction is parallel to the midrib. The other direction is perpendicular to the midrib. The number of undulations and rotation angles for each undulation are defined by user input. The frequency and amplitude of undulations are controlled by number of undulations and rotation angles, respectively. To quantify the overall degree of undulation, the average absolute rotation angles (aavg_y, aavg_z) around both axes were calculated. Since rotation angles alternate between positive and negative values to create the wave-like pattern, absolute values were used to prevent them from canceling each other out during averaging.

### 2.2 Correlation Analysis

To investigate the geometric correlations between leaf morphological parameters, a dataset of 100,000 leaf models was generated with randomized parameters. Leaf length and width were held constant at 100 cm and 8 cm, respectively, to isolate the effects of other morphological features. The maximum allowable angle for each undulation, in both the parallel and perpendicular directions to the midrib, was limited to 45°. Midrib shape parameters (M1, M2, and the right boundary of *ϕ*) were sampled from uniform distributions. Similarly, the maximum width position and the number of undulations were also uniformly distributed. The self-twisting angle followed a normal distribution with a mean of 0° and a standard deviation of 16°. Individual undulation angles, in both directions, were uniformly distributed between -45° and 45°, resulting in a normal distribution for the average absolute undulation angle. For each generated model, leaf area, projection area, and rugosity were calculated. Pearson correlation coefficients were then computed to analyze the relationships between these morphological parameters and the input parameters.

### 2.3 Optimization

This research involved two optimization tasks. The first focused on maximizing leaf area within specified morphological constraints. This involved identifying the combination of leaf shape parameters that yielded the largest possible leaf area while constrained by predefined limits on parameters such as midrib curvature, twisting, and undulation. The second optimization of canopy architecture aimed to balance leaf area and projection area to address self-shading and light distribution questions. This involved finding the optimal combination of parameters that maximized leaf area while minimizing the projection area, thereby reducing self-shading and promoting a more even distribution of light within the canopy.

#### 2.3.1 Stepwise Bayesian Optimization

To address the challenges of high-dimensional parameter optimization, a stepwise Bayesian optimization approach was adopted. This method enhances efficiency by dividing the parameter space into smaller groups, similar to the concept of optimizing via partial derivatives. After assigning initial values to all parameters, each group is optimized sequentially using a classic Bayesian optimization algorithm. This stepwise strategy is illustrated in Algorithm 1, where the parameters are partitioned into three groups, and the Bayesian optimization procedure is executed three times within a loop, with each execution targeting a specific parameter group. *N* is the user-defined iteration number. *f* is the objective function which is a black-box function and expensive to evaluate the whole feasible space. A key aspect of this method is the identification and transition between local extrema. A tolerance rate is introduced to determine when to move to the optimization of the next parameter group. Specifically, if the absolute ratio of the difference between the current parameter value and the identified local extremum, divided by the local extremum, is lower than the predefined tolerance, optimization for current group is paused. The algorithm then transitions to the optimization of the next parameter group, utilizing the optimized parameter values from the previous group as its starting point. This strategy effectively narrows the search space by focusing on localized regions around potential extrema.

**Algorithm 1.**
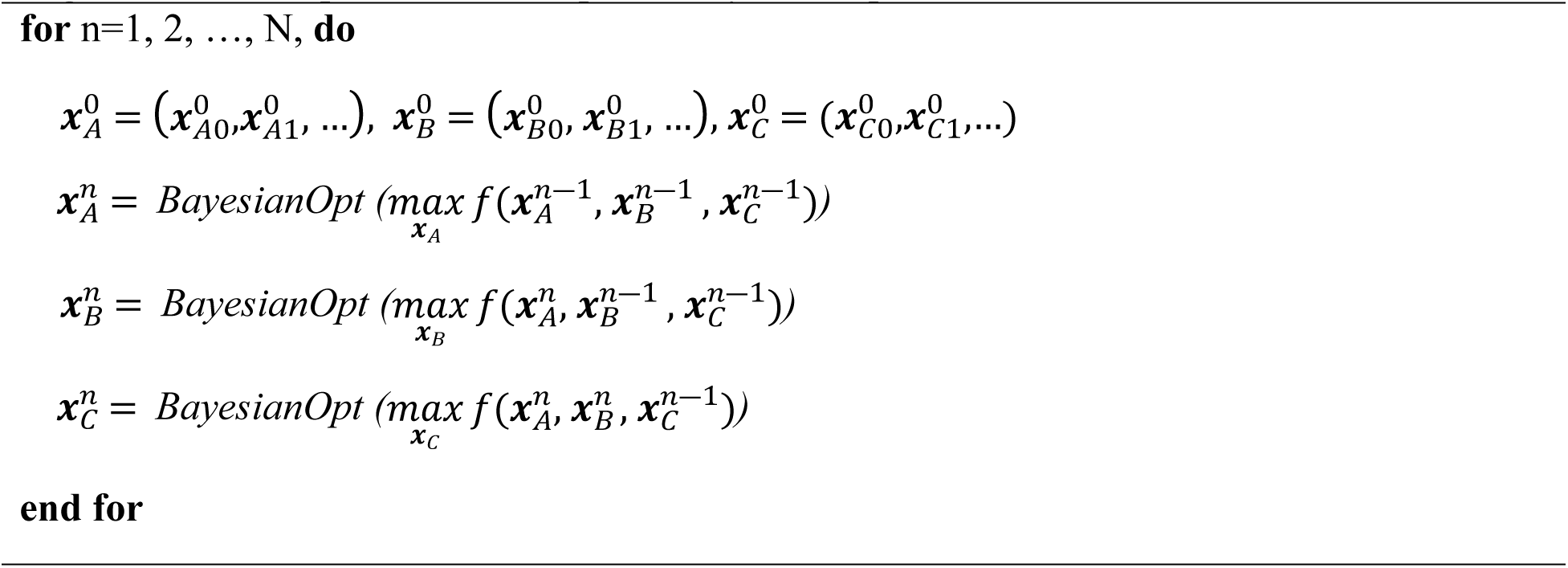
The procedure of stepwise Bayesian Optimization

#### 2.3.2 Optimization for leaf area

To optimize leaf area, a representative leaf situated in the middle canopy layer during late vegetative growth was modeled. Leaf dimensions were fixed at 100 cm in length and 8 cm in width. Given that mid-canopy leaves exhibit an intermediate midrib shape between erect upper-canopy leaves and flat lower-canopy leaves, a geometric constraint was imposed on midrib curvature. The entire midrib was constrained to lie within a shaded region bounded by three curves, as illustrated in Fig. 1a. The upper boundary (blue curve) is a hyperbolic spiral with a shape ratio of 1.4 and a right bound of 3.6, while the lower boundary (orange curve) is a hyperbolic spiral with a shape ratio of 1.8 and a right bound of 5. The right boundary (green curve) is defined by a sector of a circle with a 100 cm radius, spanning from 18° to 56°. A grid search was conducted to map the feasible region of midrib curvature, as defined by the geometric constraints of the shaded area, to the corresponding feasible space of shape ratio and right bound (Fig. 1b). The boundaries of this feasible space are defined by Eq. 2. The range for the maximum width position was set to (0.51, 0.7), and the self-twisting angle was restricted to (-10°, 10°). To investigate the impact of undulation on the leaf area, three different undulation amounts (4, 8, and 12) were considered. Undulation angles around the axes parallel and perpendicular to the midrib were limited to (0°, 30°) or (-30°, 0°), depending on the undulation position. A comparative leaf model without undulation was also included. For optimization, the morphological parameters were divided into three groups. The first group comprised midrib parameters, including shape ratio, right bound of *ϕ*, maximum width position, and self-twisting angle. The remaining two groups consisted of the rotation angles around the axes parallel and perpendicular to the midrib that generate undulations. The stepwise Bayesian method was then employed to optimize leaf area under these constraints.

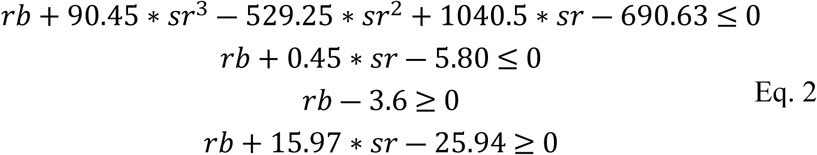

**Fig. 1.**
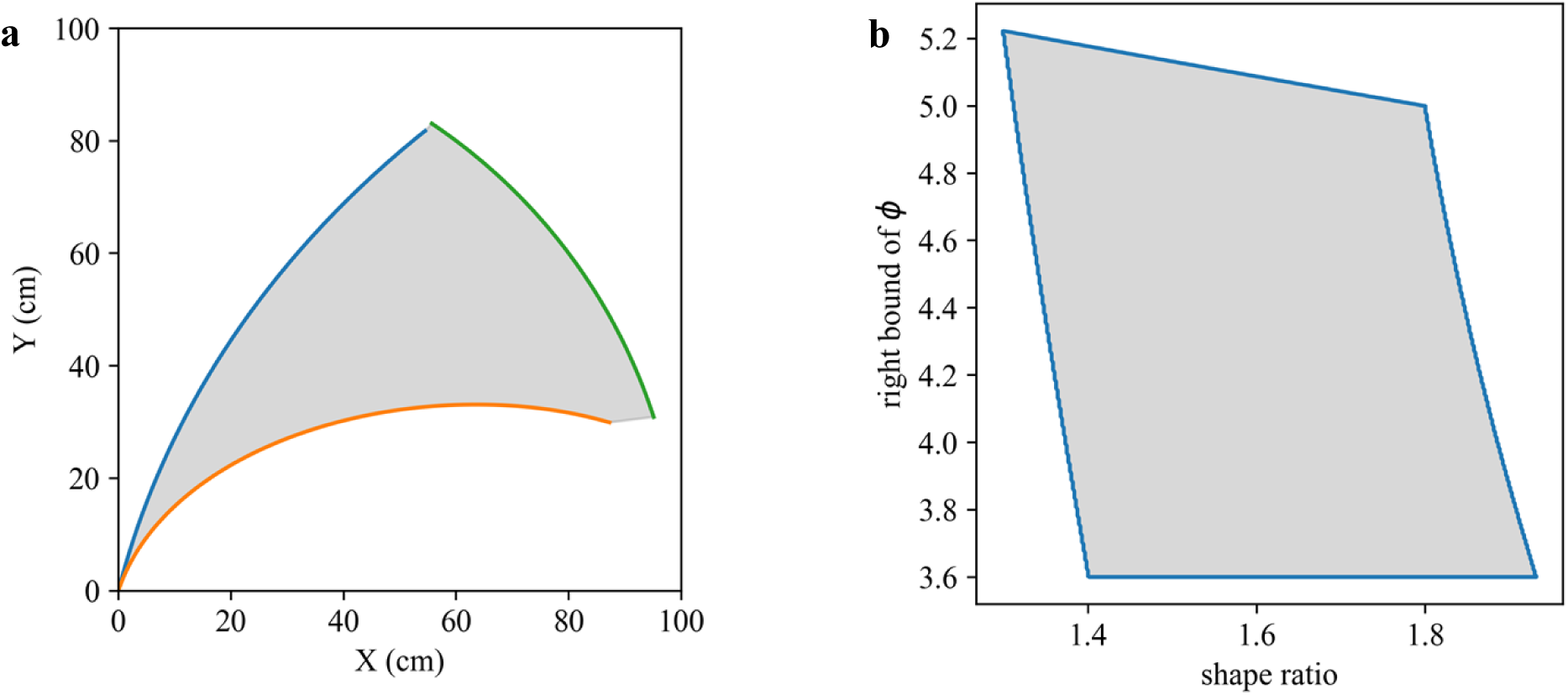
**(a)** The feasible distribution of midrib curvature. The region is enclosed by three curves. The hyperbolic spiral with a shape ratio (*sr*) of 1.4 and a right bound (*rb*) of 3.6 (blue). The hyperbolic spiral with a shape ratio of 1.8 and a right bound of 5 (orange). The sector of a circle with a 100 cm radius, spanning from 18° to 56° (green). **(b)** The feasible distribution of *sr* and *rb* according to feasible distribution of midrib curvature.

#### 2.3.3 Optimization for canopy architecture

This optimization aimed to maximize leaf area while simultaneously minimizing projection area, thereby reducing self-shading and promoting more uniform light penetration within the maize canopy. An objective function (Eq. 3) was formulated to evaluate the performance of leaf area and projection area. This function incorporates rugosity as a proxy for projection area, utilizing its ability to capture surface complexity more comprehensively. Since leaf at different layer receives inconsistent solar radiation and shading lower leaves variously, weight factor was used to adjust the standards on their performance. *w_a_* and *w_r_* are the weight factor for leaf area and rugosity, respectively. The bound was calculated based on the leaf samples in correlation analysis. The area bound (*bound_a_*) is the result of multiplication of leaf length, width and area factor. The area factor is the ratio of 95% percentile of leaf area to the maximum leaf area in the 100k leaves which is 0.87. The rugosity bound (*bound_r_*) is defined manually.

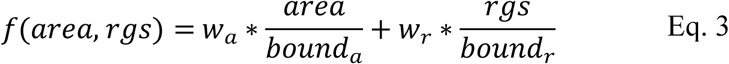

Three leaves were modeled, each representing a different canopy layer: top, middle, and bottom. The leaf at the top layer was fixed at 30 cm in leaf length and 4 cm in leaf width. Since top leaves are typically upright, a geometric constraint was imposed on the midrib curvature, requiring it to lie within the blue region depicted in Fig. 2a. This region is bounded by an upper boundary defined by a hyperbolic spiral with a shape ratio (*sr*) of 0.2 and a right bound (*rb*) of 3.5, a lower boundary defined by a hyperbolic spiral with an sr of 1.4 and an rb of 5, and a right boundary defined by a sector of a circle with a 30 cm radius, spanning from 20.2° to 85.7°. A grid search was implemented to map the feasible midrib contours to the corresponding feasible space of shape parameters (*sr* and *rb*), visualized as a blue polygon in Fig. 2b and defined by Eq. 4. Given the relatively lower surface complexity of top leaves, the undulation number was set to 4, and the undulation angle was limited to a maximum of 30°. This constraint reflects the smoother surface typically observed in top-canopy leaves compared to those in lower layers. The *w_a_* and *w_r_* were configured as 0.35 and 0.65, respectively. Because less upright top leaf intercepts more radiation and subsequently reduces the light penetration into the entire canopy. The *bound_a_* and *bound_r_* in the objective function were 104.39 and 10.45, respectively.

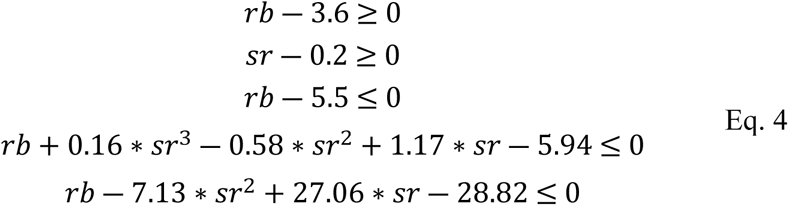

**Fig. 2.**
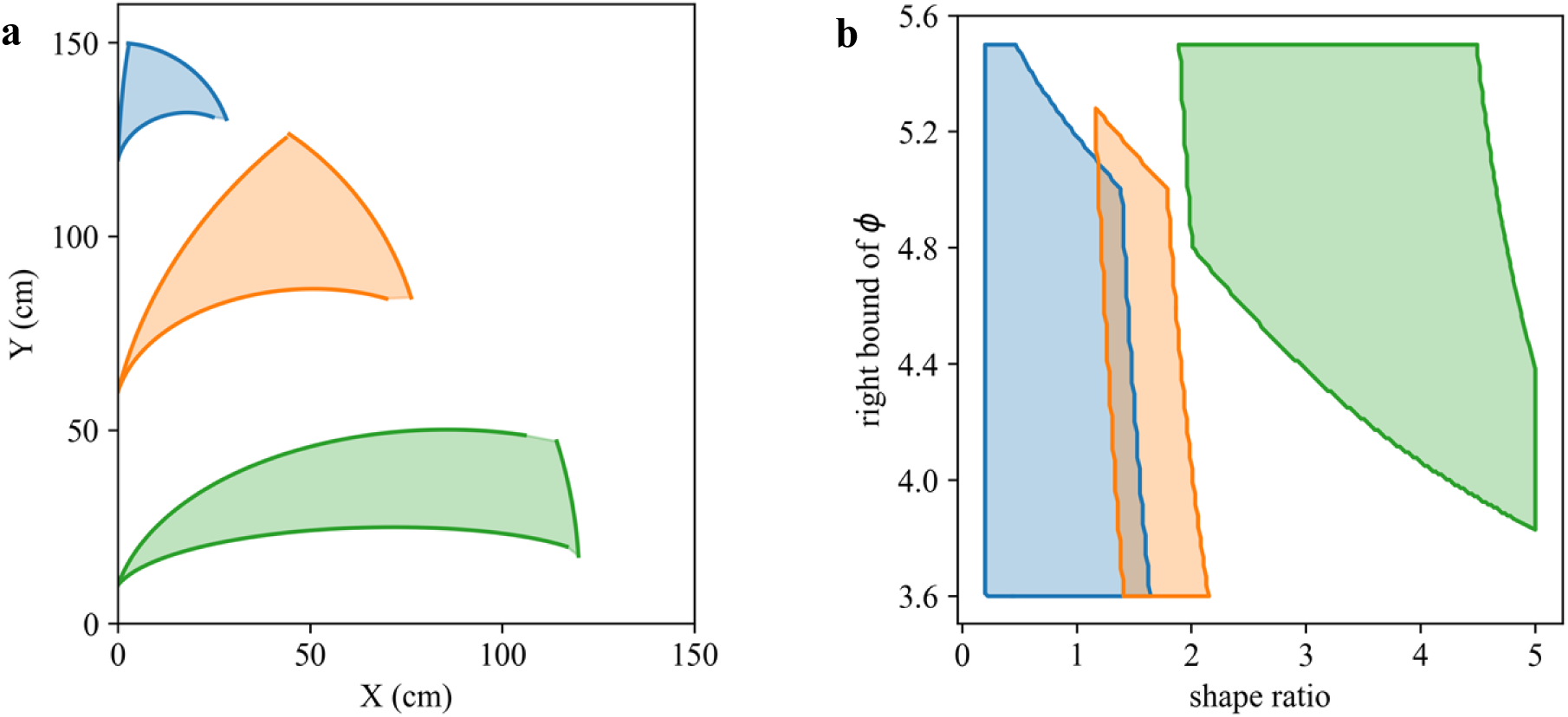
**(a)** The feasible distribution of midribs of top leaf, middle leaf and bottom leaf. The upper and lower boundaries of three leaves are hyperbolic spirals while the right boundaries are sectors of given lengths. **(b)** The feasible distribution of *sr* and *rb* of the three leaves according to their feasible curvatures.

The middle canopy leaf, with its intermediate orientation, was modeled with a fixed length of 80 cm and a width of 6 cm. To reflect this, its midrib curvature was restricted to the orange region in Fig. 2a, bounded by an upper hyperbolic spiral (*sr* = 1.4, *rb* = 3.6), a lower hyperbolic spiral (*sr* = 1.8, *rb* = 5), and a sector of a circle with radius of 80 cm, spanning from 17.6° to 56.2°. This feasible midrib space corresponds to the orange region in Fig. 2b, mathematically defined by Eq 5. With increased surface complexity compared to the top leaf, the undulation number was set to 8, and undulation angles were limited to 30°. The *w_a_* and *w_r_* were 0.5 and 0.5, respectively. The *bound_a_* and *bound_r_* were 417.55 and 1.83, respectively.

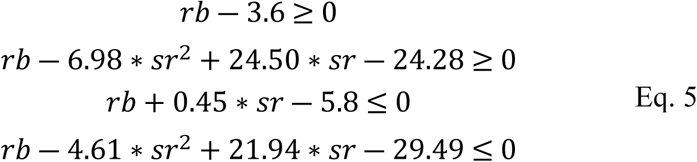

Characterized by its flatter shape and increased surface complexity, the bottom leaf was modeled with a length of 120 cm and a width of 8 cm. Undulation number was increased to 12, while maintaining the 30° limit on undulation angles. Fig. 2a depicts the feasible midrib region (green) bounded by a hyperbolic spiral with *sr* = 2 and *rb* = 4.8 (upper boundary), another hyperbolic spiral with *sr* = 4.5 and *rb* = 5.5 (lower boundary), and a sector of a circle with radius 120 cm, spanning 3.6° to 18° (right boundary). The corresponding feasible parameter space is bounded by Eq. 6, which is determined by grid search. As the bottom leaves should intercept more radiation from upper layers, the *w_a_* and *w_r_* were 0.65 and 0.35, respectively. The *bound_a_* and *bound_r_* were 835.1 and 1.13, respectively.

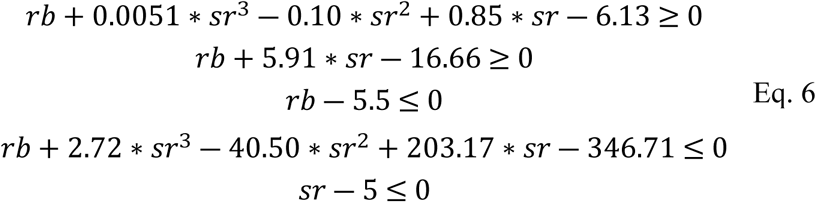

## 3. Results

### 3.1 Morphological correlations

The morphological parameters used to generate the leaf models were sampled from various probability distributions as shown in Fig. 3 with blue histogram. The shape ratio, right boundary, maximum width position, and undulation number were sampled from uniform distributions. Self-twisting angles were sampled from a normal distribution with a mean of 0° and a standard deviation of 16°. While individual undulation rotation angles were randomly generated from uniform distributions, the resulting average absolute rotation angles in both the parallel and perpendicular directions to the midrib followed a t-distribution. The distributions of undulation angles in both directions exhibited a high degree of similarity, with close means (22.53° and 22.46°) and standard deviations (5.08° and 5.13°). The degrees of freedom for the t-distributions in the parallel and perpendicular directions were 9.83 and 10.67, respectively. Leaf area, projection area, rugosity, and aspect ratio, calculated from the generated leaf models based on these input parameters, displayed more complex distribution patterns that did not conform to common distribution types. The mean leaf area and projection area were 640.29 cm² and 434.84 cm², with standard deviations of 28.95 cm² and 104.39 cm², respectively. The mean rugosity and aspect ratio were 1.65 and 0.7, with standard deviations of 0.87 and 0.88, respectively.

**Fig. 3.**
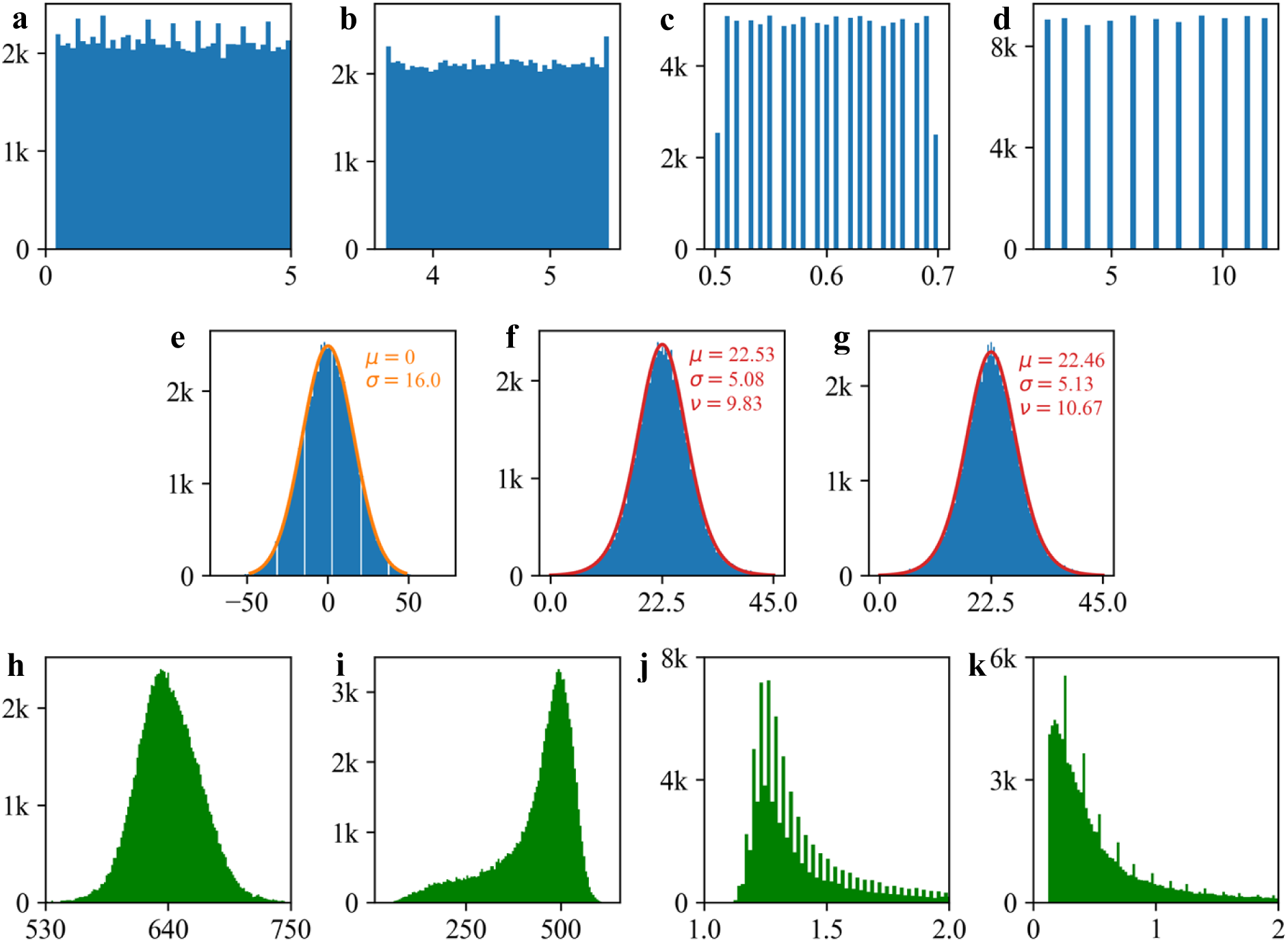
The distribution of leaf morphological parameters. The histograms with blue color are input parameters while the green are output parameters based on generated 3D leaf models. **(a)** *sr* **(b)** *rb* **(c)** *MWP* **(d)** undulation number **(e)** self-twisting angle. The orange curve represents the normal distribution with mean of 0 and standard deviation of 16. **(f)** average absolute angle of the undulations on parallel directions. The red curve is the *t*-distribution with mean of 22.53, standard deviation of 5.08 and degree of freedom of 9.83. **(g)** average absolute angle of the undulations on perpendicular directions. The red curve is the *t*-distribution with mean of 22.46, standard deviation of 5.13 and degree of freedom of 10.67. **(h)** leaf area **(i)** leaf projection area **(j)** leaf rugosity **(k)** leaf aspect ratio

The correlation analysis reveals several key relationships between leaf morphology variables. The result of correlation is depicted in Fig. 4. Rugosity and aspect ratio exhibit a very strong positive correlation (0.968), indicating that leaf surface complexity is closely tied to midrib curvature. Projection area is strongly negatively correlated with both rugosity (-0.862) and aspect ratio (- 0.882), suggesting that leaves with larger projection areas tend to be simpler in shape. The midrib shape ratio has moderate correlations with several variables, influencing leaf area (-0.319), projection area (0.775), rugosity (-0.603), and aspect ratio (-0.633). In contrast, undulation number and self-twisting show weak linear correlations with other parameters. Other notable correlations include moderate positive correlations between leaf area and the right boundary parameter (0.454), and between leaf area and maximum width position (0.361). Most of the remaining correlations are weak, indicating little to no strong linear relationship between those variable pairs. Undulation has a small positive correlation with leaf area (0.132), suggesting that leaves with more undulations tend to have a slightly larger surface area. However, the undulation angles in parallel and perpendicular directions display divergent effects. The undulation angles in the parallel direction show a negligible positive correlation with leaf area (0.05), while the undulation angles in the perpendicular direction show a moderate negative correlation with leaf area (-0.37). The strong correlations among rugosity, aspect ratio, and projection area suggest these variables are central to leaf shape and its interaction with light, where high rugosity and aspect ratio (complex, curved shapes) are associated with smaller projection areas (less shading), likely reflecting a trade-off between maximizing light capture and minimizing shading of lower leaves.

**Fig. 4.**
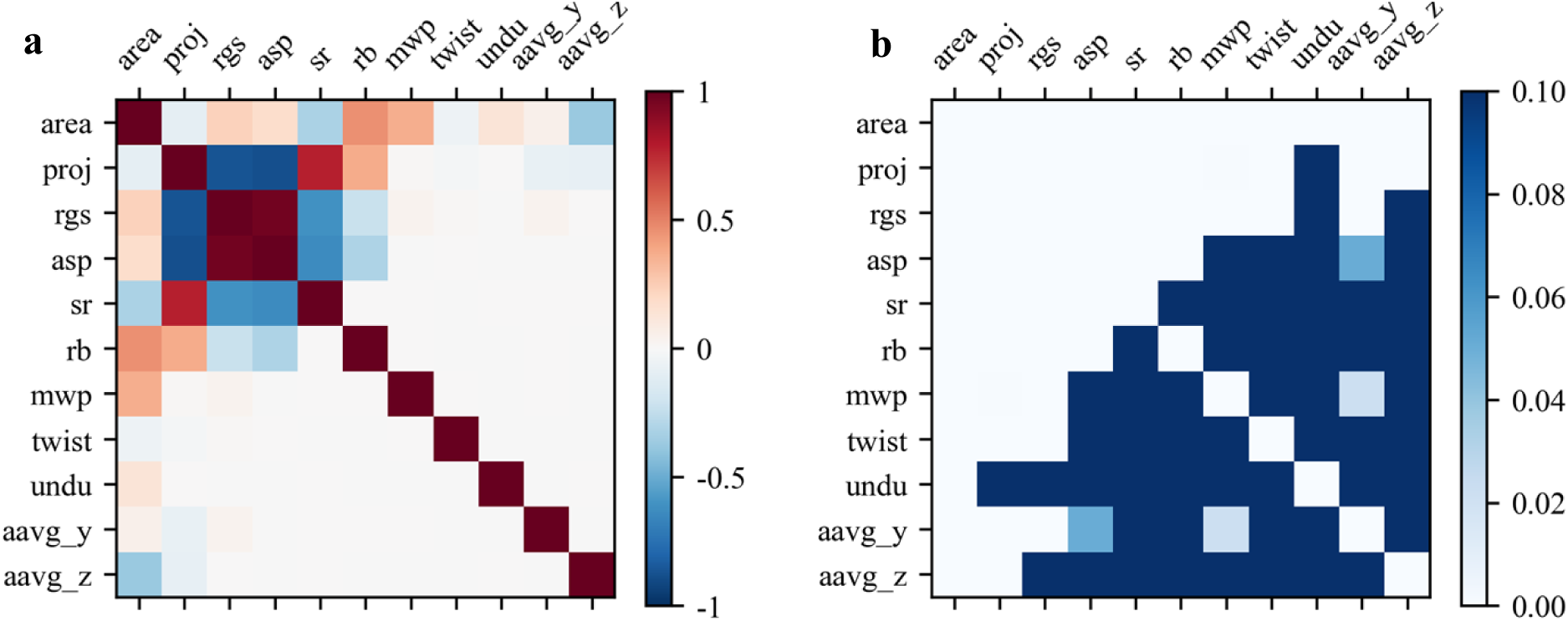
The correlation (**a**) and *p*-value (**b**) for morphological parameters, including leaf area (area), leaf projection area (proj), rugosity (rgs), aspect ratio (asp), shape ratio of midrib (sr), right boundary of the intermediate parameter *ϕ* (rb), maximum width position (mwp), self-twisting (twist), undulation number (undu), average absolute undulation angle in parallel direction (aavg_y), average absolute undulation angle in perpendicular direction (aavg_z).

### 3.2 The impacts of undulation amount and position

The amount and position of undulation significantly influence key leaf morphology parameters, including leaf area, projection area, and rugosity. Correlation analysis, as shown in Fig. 5, reveals that undulation angles in the parallel direction to the midrib tend to marginally increase leaf area. This suggests that parallel undulations might expand the leaf surface in a way that contributes to a slight increase in overall area. However, undulation angles in the perpendicular direction to the midrib tend to slightly decrease leaf area. This indicates that perpendicular undulations compress the leaf surface, leading to a reduction in surface area as the leaf is effectively “folded” in that direction. Undulation angles in both directions marginally reduce leaf projection area. This implies that undulations, regardless of their orientation, tend to create more complex shapes that cast smaller shadows compared to a flat leaf. Consequently, undulations contribute to increased leaf surface complexity, resulting in higher rugosity. The increased surface area relative to the projected area signifies a more convoluted leaf structure. Furthermore, the position of undulations affects the extent of their impact on leaf area. Undulations located in the middle region of the leaf, where leaf width is greater than at the leaf tip, have a greater impact on leaf area. This suggests that the broader surface in the mid-region provides more material to be affected by the undulations, leading to more pronounced changes in the overall leaf area.

**Fig. 5.**
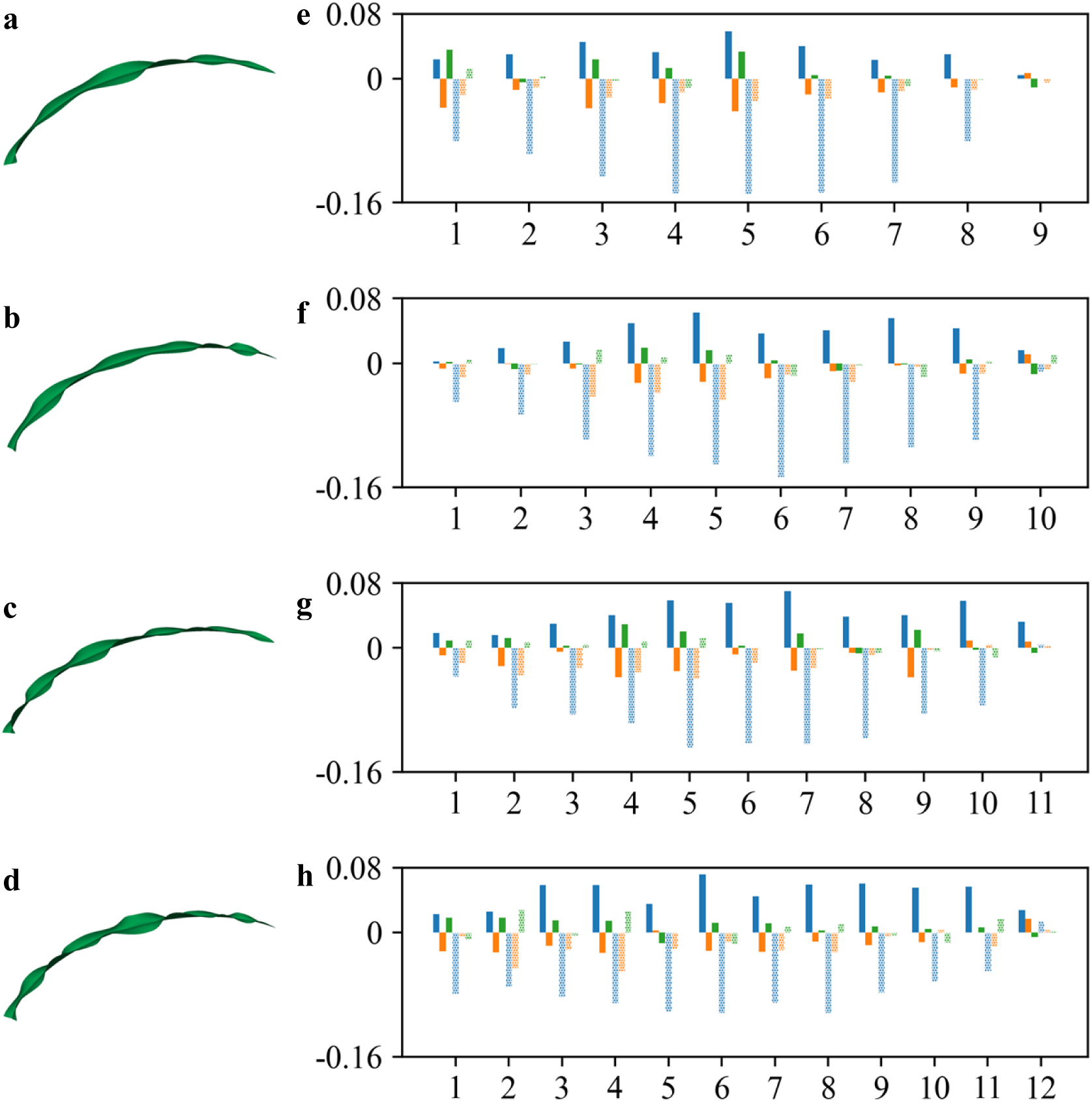
The impacts of undulation amount and position. **(a-d)** Leaf models with 9, 10, 11, 12 undulations. **(e-h)** The correlation between undulation positions and leaf area (blue), projection area (orange) and rugosity (green) in both parallel (solid bar) and perpendicular (dotted bar) directions for leaves with 9, 10, 11, 12 undulations.

### 3.3 Optimization of leaf area

The optimization of leaf morphology, subject to geometric constraints and conducted over 100 iterations, revealed distinct trends in leaf area and other key properties across four undulation conditions (0, 4, 8, and 12 undulations). The optimization trajectory and result are shown in Fig. 6. Leaf area steadily increased, ultimately converging to maximum values of 704.196 cm², 704.884 cm², 708.328 cm², and 711.817 cm² for the respective undulation conditions. This demonstrates a clear relationship between increased undulation and larger leaf area. Concurrently, projection area exhibited initial variability but ultimately stabilized around 512.412 cm², 487.594 cm², 483.193 cm², and 486.235 cm² for the corresponding undulation conditions. The ratio of leaf area to projection area, representing rugosity, converged to 1.374, 1.446, 1.466, and 1.464, respectively, indicating that increased undulation leads to greater surface complexity. The aspect ratio, primarily determined by midrib shape, converged to similar values across all undulation conditions (0.481, 0.480, 0.482, and 0.473), reflecting the proximity of the optimized midrib parameters. The shape ratio (1.301, 1.304, 1.306, 1.332), and right boundary of *ϕ* (5.214, 5.123, 5.199, 5.196) showed only minor variations. The maximum width position (0.666, 0.670, 0.667, 0.676) and self-twisting (5.492°, 6.184°, 7.569°, 8.773°) exhibited oscillated patterns. Furthermore, leaf self-twisting angles increased with undulation number. The average absolute undulation angles demonstrated a trend of increasing complexity, with values of 0°, 24.2°, 22.94°, and 23.72° for the parallel direction and 0°, 0°, 4.33°, and 8.54° for the perpendicular direction. Notably, 0° indicates the absence of undulation in that direction.

**Fig. 6.**
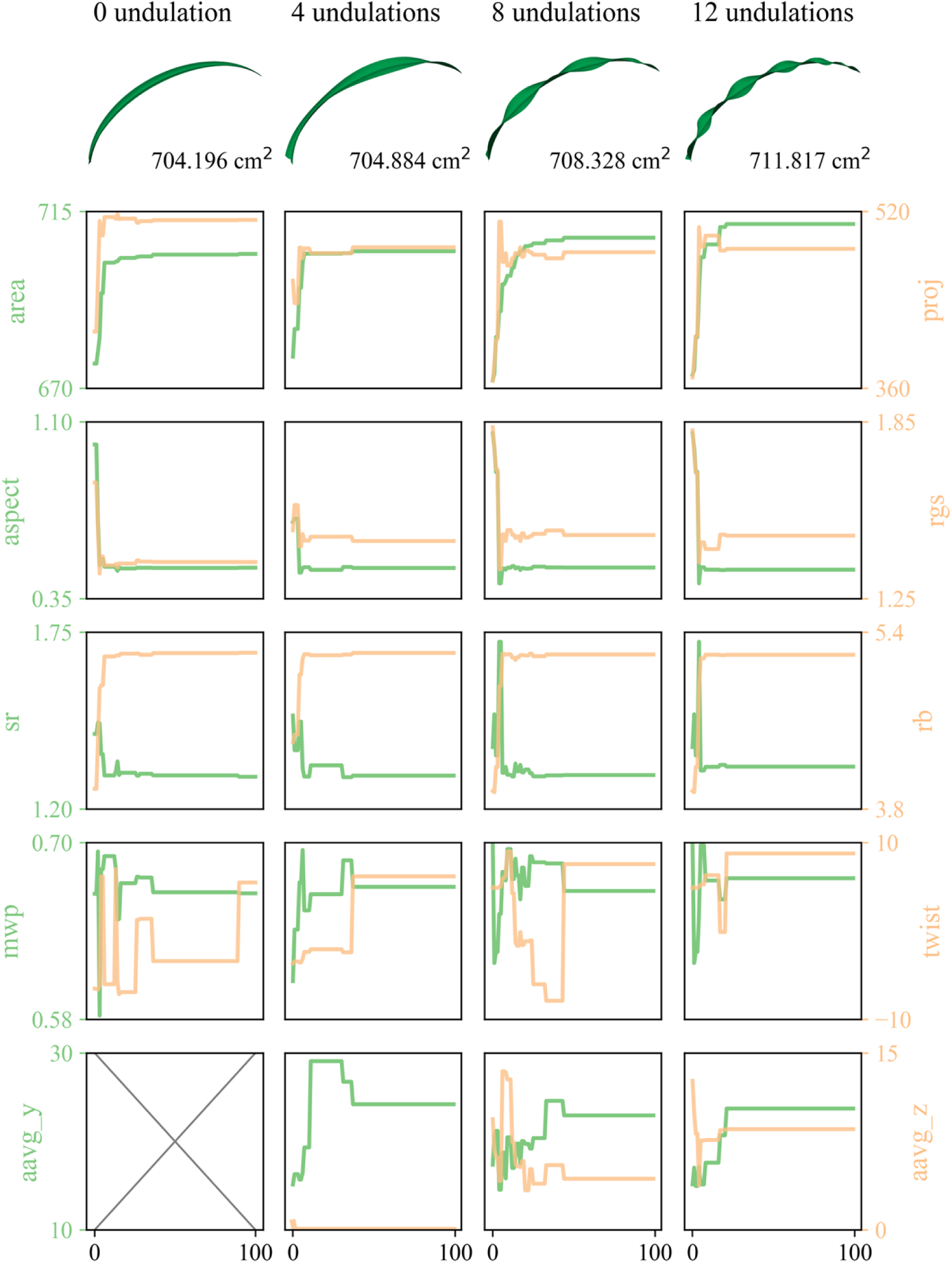
The stepwise Bayesian optimization trajectory and results for leaf area with 0, 4, 8 and 12 undulations under the constraints of leaf length and width.

### 3.4 Optimization of rugosity and leaf area

The optimization of leaf area and projection area for canopy architecture was performed over 100 iterations using a customized objective function. This function incorporated leaf area and rugosity, with varying weights assigned to each parameter depending on the leaf’s position within the canopy (top, middle, or bottom). This weighting scheme allowed for a standardized comparison of leaves with different dimensions and properties. Initial leaf lengths were set to 30 cm, 80 cm, and 120 cm for the top, middle, and bottom leaves, respectively, with corresponding widths of 4 cm, 6 cm, and 8 cm. As depicted in Fig. 7, the optimization process yielded distinct results for each canopy layer. The objective function values for the top, middle, and bottom leaves converged to 1.24, 1.13, and 1.06, respectively. Notably, the top and middle leaves exhibited a greater increase in their objective function values compared to the bottom leaf. This can be attributed to the emphasis on minimizing self-shading for the upper canopy layers. The top leaf displayed oscillations in leaf area before converging at 109.47 cm² that was near the initial values of 111.81 cm². Its projection area consistently decreased from 10.38 cm² to 7.79 cm², resulting in a substantial increase in rugosity from 10.77 to 14.05. The aspect ratio initially increased from 9.80 to 10.48 before settling at 10.43. These values were significantly higher than those observed for the middle (aspect ratio: 1.48, rugosity: 2.39) and bottom (aspect ratio: 0.3, rugosity: 1.33) leaves. The middle leaf’s area oscillated before reaching a final value of 401.41 cm², while its projection area generally decreased from 273.32 cm² to 167.87 cm². The bottom leaf, with the largest dimensions, achieved a final leaf area of 839.73 cm² and a projection area of 630.82 cm². The optimized midrib shape parameters also varied across the canopy layers. The top leaf had the smallest shape ratio (0.2), compared to 1.41 for the middle leaf and 1.9 for the bottom leaf. The right boundary of *ϕ* was 3.6 for both the top and middle leaves, increasing to 5.5 for the bottom leaf. Maximum width positions were 0.62, 0.7, and 0.67 for the top, middle, and bottom leaves, respectively. Self-twisting angles were -9.04°, 8.28°, and 3.09° for the top, middle, and bottom leaves, respectively. Average absolute undulation angles in the parallel direction were 11.83°, 14.94°, and 20.97°, while those in the perpendicular direction were 14.33°, 18.07°, and 16.64°. These angles exhibited oscillations throughout the optimization process.

**Fig. 7.**
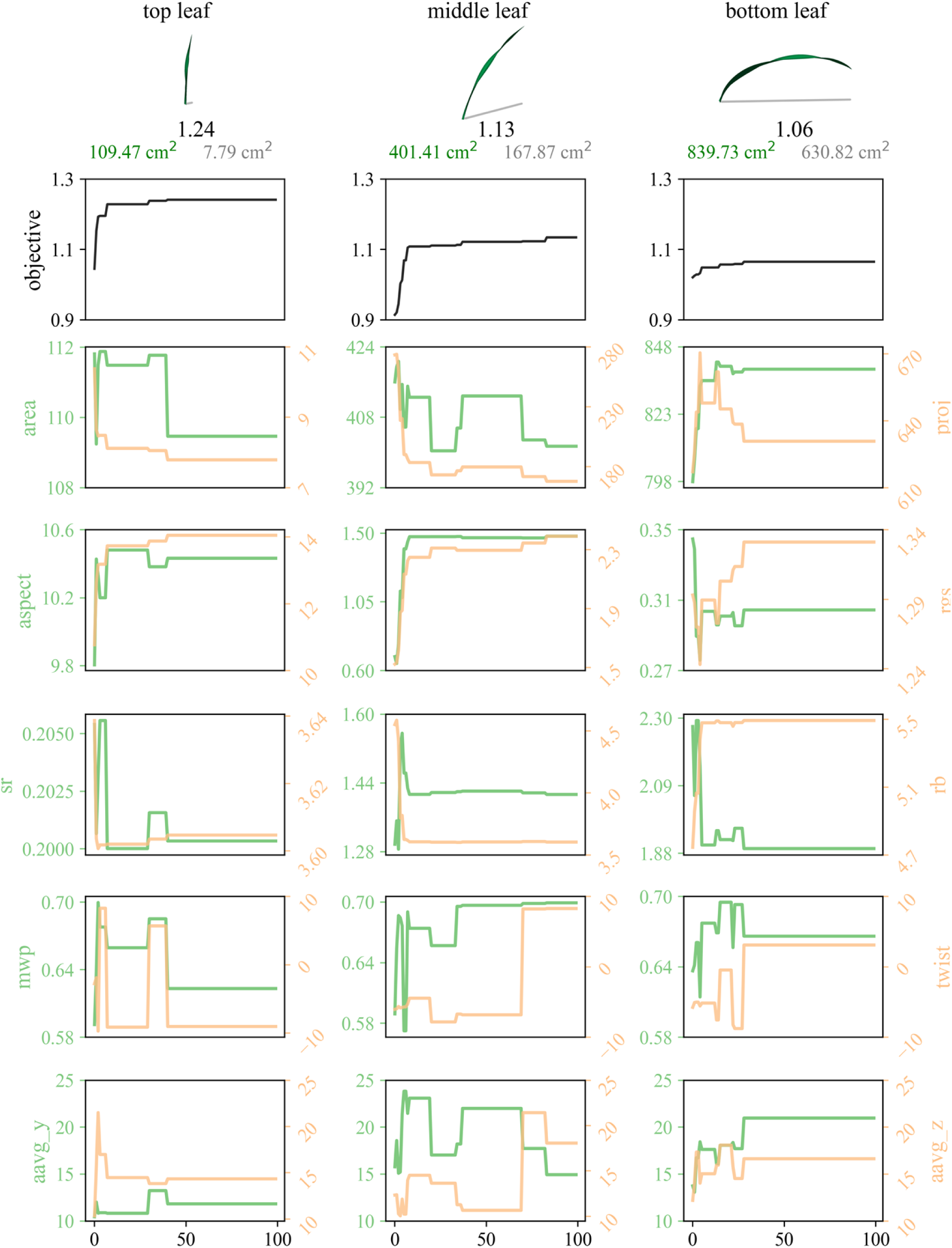
The stepwise Bayesian optimization trajectory and results for canopy architecture under constraints of midrib shapes, leaf length and width over 100 iterations.

## 4. Discussion

### 4.1 Leaf Development and Its Morphology

The maize leaf is composed of three primary structural components: the blade, the sheath, and the auricle/ligule. Leaves are exposed to a variety of environmental factors, including light, water, temperature, microbes, and insects, and serve as an important medium for environmental interaction. The leaf is the primary organ for photosynthesis, the process by which carbon dioxide and water are converted into carbohydrates and oxygen, which is essential for plant growth and development. Our research investigates the morphology of the maize leaf blade, the expanded portion of the leaf responsible for the majority of light capture. Understanding leaf blade morphology is of paramount importance as it serves as a key determinant of photosynthetic efficiency, directly impacting the plant’s overall productivity and yield. Furthermore, Leaf morphology effectively reflects maize growth and health status, providing valuable insights into the plant’s physiological condition. However, leaf morphology is dynamic, exhibiting significant variability even under relatively constant conditions, such as those maintained within a greenhouse environment. Furthermore, leaf morphology demonstrates high responsiveness to environmental fluctuations due to plant plasticity, including variations in light intensity, temperature, and water availability. The intricate mechanisms governing the attainment of final maize leaf morphology are not fully understood. These unresolved questions include the precise molecular and cellular processes underlying leaf initiation, the complex genetic and hormonal networks that control leaf shape determination, and the establishment and maintenance of leaf polarity (Strable & Nelissen, 2021). Leaf shape is dominantly determined by leaf polarity, as differential growth patterns and cell differentiation along the leaf axes establish the diverse leaf morphologies. Leaf polarity is a fundamental aspect of leaf development, established and maintained during growth, encompassing three primary axes: adaxial-abaxial, proximal-distal, and medio-lateral. The regulatory network controlling leaf polarity is complex and involves a multitude of interacting genes and signaling pathways. Adaxial-abaxial polarity is initiated very early in development, even before leaf initiation (Husbands et al., 2009). This process involves distinct sets of genes controlling the development of adaxial and abaxial cell fates. For example, the *REVOLUTA* (*REV*) gene, encoding an HD-ZIP III transcription factor, plays a critical role in defining adaxial cell fate (Emery et al., 2003). After leaf initiation, cells expressing the *KANADI1* (*KAN1*) gene in the shoot apical meristem are specified to adopt an abaxial identity (Kerstetter et al., 2001). *YABBY* genes also contribute to the establishment and maintenance of dorsoventral polarity (Juarez et al., 2004). The distal blade and proximal sheath, two functionally and morphologically distinct tissues, differentiate in a process dominantly regulated by *knotted-like homeobox* (*knox*) mutants, a family of transcription factors involved in regulating cell fate and differentiation (Ramirez et al., 2009). The *ragged seedling2* (*rgd2*) gene, encoding an ARGONAUTE7-like protein, is required for proper medio-lateral expansion, controlling leaf width and overall blade size (Douglas et al., 2010). The *drooping leaf1* (*drl1*) *and drooping leaf2* (*drl2*) genes regulate leaf support tissues and restrict auricle expansion at the midrib (Strable et al., 2017). In maize expressing *drl1* and *drl2*, leaves exhibit reduced erectness, drooping at the proximal midrib. Despite these significant advances in our understanding of leaf development, a comprehensive and integrated understanding of the genetic regulatory network governing the intricate processes of leaf growth and development remains elusive. One of the major factors contributing to this complexity is the spatial variation of cell types and its gene expression along the maize leaf. A leaf in the vegetative stage can be divided into four distinct regions along its proximal-distal axis from the base to the tip: the basal zone, the transitional zone, the maturing zone, and the mature zone (Li et al., 2010). These four zones exhibit distinct patterns of growth and morphology, reflecting their specialized functions. Furthermore, the gradient of gene expression decreases from base to tip, reflecting the complex reprogramming mechanisms associated with the gradual development of photosynthetic function as the leaf matures. Thus, the leaf transitions from a base focused on fundamental growth and development, including cell division and differentiation, through a middle zone establishing the photosynthetic machinery, including chloroplast development and chlorophyll biosynthesis, to a tip optimized for efficient photosynthesis, characterized by high levels of Calvin cycle enzymes and photosystem components.

### 4.2 Leaf Morphology and Canopy Architecture

Maize canopies exhibit high geometric complexity. While leaf area index (LAI) and light extinction coefficient (*k*) are commonly employed to describe canopy architecture, these indicators are primarily designed to address radiation-related performance. Canopy architecture is fundamentally determined by leaf number and leaf morphology. However, descriptions of maize canopies based solely on leaf morphology are often insufficient to capture this complexity. Traditionally, among the many leaf morphological traits, leaf area, leaf elevation angle, and azimuth angle have been considered dominant factors influencing canopy architecture and light interception ability. Further research on the intricate relationship between leaf area and canopy architecture has been limited by the inherent complexity of canopy geometry and the labor-intensive nature of field work required to collect detailed leaf and canopy information.

Over the past decades, breeding efforts in major maize planting regions have favored increased leaf erectness and more compact canopies to facilitate higher planting densities and enhance yield (Messina et al., 2009; Perez et al., 2019; Sangoi et al., 2002; Zhao et al., 2015). This erectophile leaf habit promotes more even vertical distribution of light and reduces *k* within the canopy. However, optimal radiation use efficiency likely requires distinct leaf morphological patterns at different canopy layers. Indeed, an optimized canopy architecture often features upright upper leaves, less erect middle leaves, and relatively flat lower leaves (Tian et al., 2019, 2024). Leaf angle, particularly the elevation angle, is a major determinant of canopy architecture. The leaf angle is influenced by the structural characteristics of the region between the sheath and blade, including the ligule and auricle. Consequently, genes regulating ligule and auricle development play a significant role in controlling leaf elevation angle. The *liguleless1* (*lg1*) mutant lacks both ligule and auricle, resulting in a compact canopy architecture with reduced leaf angle (Sylvester et al., 1990). In *liguleless2* (*lg2*) plants, the ligule and auricle are absent or mispositioned, and the blade-sheath boundary is diffuse (Walsh et al., 1998). Other genes in the *KNOX* family, including *liguleless3* (*lg3*), *liguleless4* (*lg4*), *knotted* (*kn1*), and *rough sheath1* (*rs1*), also regulate ligule/auricle regions directly (Jiang & Wang, 2025). Some genes regulate leaf angle indirectly through phytohormones, such as brassinosteroids (BR) and gibberellins (GA). *Brassinosteroid insensitive1* (*bri1*) regulates phytohormone signaling and consequently affects ligule and auricle development (Kir et al., 2015). It encodes a leucine-rich repeat receptor-like kinase and plays a vital role in brassinosteroid signaling pathways. The *nana plant2* (*na2*) mutant exhibits decreased levels of downstream BR metabolites and possesses a compact architecture with reduced leaf angle (Best et al., 2016). While leaf elevation angle has received considerable attention, the leaf azimuth angle also contributes to canopy architecture. Research on leaf azimuth angle is relatively limited. Azimuth angle distribution is regulated by plant phototropism. *liguleless* genes result in low auxin levels at the leaf base, negatively affecting the plant’s phototropic response and thus influencing azimuth angle (Zhou et al., 2024). Canopy architecture is not solely determined by leaf angle. Midrib shape and internode length also contribute to its complexity. However, the impacts of these other leaf morphological properties on canopy architecture have received less attention from breeders and geneticists. Addressing this gap is crucial for maize ideotype design and breeding for radiation use efficiency (Huang et al., 2024; Jafari et al., 2024).

Our optimization of canopy architecture provides a quantitative framework to evaluate the performance of a given canopy geometry, with a focus on increasing leaf area and decreasing shadow projection below. Although this framework represents a simplified description of the three-dimensional architecture of maize canopies, which encompasses a multitude of interacting factors such as leaf curvature, blade surface and leaf position, it offers a valuable and essential tool for quantifying the inherent complexity of leaf and canopy geometry. This quantification, by providing precise metrics for previously subjective traits, allows for more rigorous analysis of how these geometric features vary and interact. This capability is crucial for advancing quantitative genetic studies, enabling researchers to accurately correlate genetic variation with the observed complex geometric variation in leaf and canopy structure.

### 4.3 Model-Assisted Plant Phenotyping

Plant phenotyping has become a widely utilized approach for monitoring plant growth and development across various crops, growth stages, platforms, and scales (Yang et al., 2020). Plant morphology is a primary focus of plant phenotyping efforts. However, significant challenges remain in the effective detection and measurement of plant morphology. These challenges stem from two interwoven factors: the inherent morphological complexity of plants, which is both time-variant and shape-dynamic, and the lack of robust quantitative methods for characterizing these complex traits. For example, the shape and structure of maize ligule and auricle region, which play a crucial role in leaf morphology by supporting the leaf blade and experiencing high torque, are still inadequately understood (Edson-Chaves et al., 2023). Developing quantitative and biophysical models for this region would enable the extraction of torque information and facilitate separate analysis of midrib and blade properties.

Functional-structural plant models (FSPMs) offer a promising framework for integrating plant morphology and physiology (Louarn & Song, 2020). FSPMs can incorporate numerous quantified morphological traits to model the complex interactions between environment, management practices, and genotype. Moreover, FSPMs can potentially reduce the frequency of data collection required for monitoring growth throughout the season by providing interpolation based on programmed growth patterns. Furthermore, while current plant phenotyping often focuses on basic traits such as leaf length, leaf angle, and biomass, these traits may not comprehensively and accurately represent plant growth and health. For instance, water use efficiency (WUE), an important indicator of irrigation effectiveness, cannot be directly measured or detected by current phenotyping methods (Hatfield & Dold, 2019). However, plant features affecting WUE, including early vigor, root architecture, canopy temperature depression, and osmotic adjustment, are related to basic traits. Examples include seedling growth rate and leaf area ratio for early vigor, and maximum rooting depth and root distribution index for root architecture. Plant phenotyping primarily addresses the detection of these basic traits. A gap remains between these basic traits and the comprehensive assessment of plant health and performance. Therefore, a systematic phenotyping approach integrated with FSPMs is necessary to bridge this gap.

### 4.4 Limitations

First, the representation of leaf undulation in our model is a simplification of the complex curvature and surface characteristics observed in real leaves. While our model employs an equidistant distribution of undulations across the leaf blade, the actual distribution is likely more complex and diverse. A significant challenge in accurately modeling leaf undulation is the current lack of robust methods for its precise measurement and detection, which limits the availability of reliable reference data for model development and validation. Second, our model’s representation of the leaf blade cross-section using a tractrix curve may not fully capture the complexity of actual maize leaf cross-sections. Although the tractrix curve’s shape can be adjusted by parameters, maize leaves may exhibit more intricate cross-sectional patterns influenced by drought stress, nutrient availability, and disease. Third, the numerical and geometrical description of canopy architecture in this study is simplified compared to the complicated reality of natural canopies. The inherent complexity of three-dimensional canopy structure is challenging to represent and interpret using two-dimensional techniques. Future studies could benefit from the application of three-dimensional imaging technologies, such as LiDAR and depth cameras, to enable more comprehensive analysis of leaf morphology and canopy architecture.

## 5. Conclusion

This study explored the relationships between midrib shape, blade contour, undulations, and surface complexity in maize leaves. We found that midrib shape and leaf width distribution positively influence leaf area. Undulation has a bidirectional effect on leaf area: undulation angles parallel to the midrib tend to increase leaf area, while perpendicular undulation angles tend to decrease it. Midrib shape and undulation also contribute to surface complexity, with increased steepness and erectness enhancing complexity by reducing projection area. Furthermore, undulation position affects leaf area and surface complexity, with greater effects observed for undulations located in regions of greater leaf width. We employed a stepwise Bayesian optimization method to identify optimal leaf morphologies for leaf area and canopy architecture.

The optimization results indicate that leaves with the largest area tend to have relatively flat midribs and a maximum width located near two-thirds of the leaf length. The optimization also revealed that the effect of undulation amount on leaf area is marginal. Optimization of canopy architecture, aimed at maximizing light radiation distribution within the canopy, considered both leaf area and shadow (represented by projection area) in the objective function. The results demonstrate that optimal canopy architecture features upright top leaves, intermediately erect middle leaves, and relatively flat bottom leaves, with decreasing midrib curvature from top to bottom leaves. In summary, this study successfully identified leaf morphologies that balance the competing objectives of maximizing leaf area and minimizing self-shading, resulting in distinct architectural characteristics for each canopy layer. This provides a quantitative framework for understanding and predicting optimal leaf morphology and canopy architecture.

## Acknowledgements

This research is supported by USDA-NIFA grants titled “High Intensity Phenotyping Sites: Transitioning to a nationwide plant phenotyping network”.

